# A Parallel Multiobjective Metaheuristic for Multiple Sequence Alignment

**DOI:** 10.1101/103101

**Authors:** Álvaro Rubio-Largo, Leonardo Vanneschi, Mauro Castelli, Miguel A. Vega-Rodríguez

## Abstract

The alignment among three or more nucleotides/amino-acids sequences at the same time is known as Multiple Sequence Alignment (MSA), an NP-hard optimization problem. The time complexity of finding an optimal alignment raises exponentially when the number of sequences to align increases. In this work, we deal with a multiobjective version of the MSA problem where the goal is to simultaneously optimize the accuracy and conservation of the alignment. A parallel version of the Hybrid Multiobjective Memetic Metaheuristics for Multiple Sequence Alignment is proposed. In order to evaluate the parallel performance of our proposal, we have selected a pull of datasets with different number of sequences (up to 1000 sequences) and study its parallel performance against other well-known parallel metaheuristics published in the literature, such as MSAProbs, T-Coffee, Clustal Ω, and MAFFT. The comparative study reveals that our parallel aligner is around 25 times faster than the sequential version with 32 cores, obtaining a parallel efficiency around 80%.

## 1 Introduction

Multiple Sequence Alignment (MSA) is the process of aligning three or more nucleotides/amino-acids sequences at the same time [1]. The main aim of MSA is to discover common ancestors among biological sequences. The discovery of biological relationship among several sequences is vital for inferring phylogenetic relationships among groups of organisms [4] [7]. Another important goal of MSA is the determination of biological significance among the given sequences [14], therefore, we prioritize the conservation within regions throughout the alignment process. Finally, a proper alignment helps us to detect which regions of a gene are susceptible to mutation and which can have one residue replaced by another without changing the function.

The MSA problem is an NP-complete optimization problem; therefore, different heuristic approaches have been developed for solving this problem in an effcient amount of time. In the literature, we find three main groups: *progressive* methods, *consistency-based* methods, and *iterative refinement* methods.

The most representative *progressive* methods are: Clustal W [20], Clustal Ω [19], PRANK [13], and Kalign [11]. These methods start calculating a distance matrix from every pair of the given sequences; then, a guide tree is built by using any hierarchical clustering algorithm, such as Unweighted Pair Group Method with Arithmetic Mean (UPGMA); finally, the alignment is obtained by following the guide tree. The main disadvantage of *progressive* methods relies on the chance of including an inaccurate gap at the beginning that will be propagated to final alignment.

The second group includes those methods based on *consistency*. These approaches construct a database of local and global alignments between each pair of sequences that helps to build an accurate multiple alignment among all the given sequences. Among the most important consistency-based tools are: Tree-based Consistency Objective Function For alignment Evaluation (T-Coffee) [15], PROBabilistic CONSistency-based multiple sequence alignment (ProbCons) [3], and MSAProbs [12].

Finally, we find the *iterative refinement* tools. The methodology followed by these tools starts by performing a progressive alignment and then, they iterate with the aim of correcting any possible inaccurate gap inserted in the progressive construction stage. The refinement process is repeated until no further improvements are found or until a predefined number of iterations is reached. Among the most widely-used *iterative refinement* methods, we find: MUltiple Sequence Comparison by Log-Expectation (MUSCLE) [5] and Multiple Alignment using Fast Fourier Transform (MAFFT) [10].

In this paper, we propose the use of Multiobjective Optimization and Evolutionary Computation in order to optimize simultaneously quality and consistency of the final alignment. We have already applied these techniques to other optimization problems in the Bioinformatics [8] and Telecommunication fields [17], [16]. A memetic metaheuristic has been chosen for this purpose, the Shuffled Frog-Leaping optimization Algorithm (SFLA) [6], which is based on the evolution of memes carried by the interactive individuals, and a global exchange of information among themselves. The traditional SFLA has been modified for optimizing multiple objective functions simultaneously and hybridized with a local search procedure. We refer to it as a Hybrid Multiobjective Memetic Metaheuristic for the Multiple Sequence Alignment (H4MSA).

Given a set of k non-aligned sequences where the length of the largest sequence is L, the time and space complexity for solving the MSA problem is O(*k*2^*k*^L^*k*^) [22]; therefore, to find an optimal alignment of a large number of sequences (~1000 sequences) becomes computationally intractable. Some of the aforementioned tools allow parallelism: Clustal Ω, T-Coffee, MSAProbs, and MAFFT; however, it is still a challenge to develop accurate methods that provide higher parallel efficiencies for aligning very large sets of sequences. The main contribution of this paper is an efficient parallel version of H4MSA for aligning very large sets of sequences accurately.

The rest of the paper is organized as follows. Section 2 formulates the Multiple Sequence Alignment problem. A detailed description of the parallel H4MSA algorithm is presented in Section 3. Section 4 is devoted to compare the alignment accuracy and parallel efficiency of H4MSA with other parallel approaches published in the literature. Finally, the conclusions and future works are presented in Section 5.

## 2 Multiple Sequence Alignment Problem

Given a set of sequences *S*: {*s*_1_, *s*_2_, … *s_k_*} of lengths |*s*_1_|, |*s*_2_|, …, |*s_k_*| defined over an alphabet Σ, such as Σ = *_nucleotides_* = {*A, C, G, T*} or Σ = *_aminoacids_* = {*A, C, D, E, F, G, H, I, K, L, M, N, P, Q, R, S, T, V, W, Y*}.

A multiple sequence alignment of S is defined as *S*^′^: {*s*^′^_1_, *s*^′^_2_, …, *s*^′^_*k*_}, where the length of the k sequences is exactly the same. Note that, S^′^ is defined over the same alphabet as *S* (*Σ*) with an additional gap symbol (−); so, *S*^′^ is defined over the alphabet Σ ∪ [{−}.

In this way, a multiple alignment is obtained by adding gaps to the sequences of *S* so that their lengths become the same. It can be seen as a matrix representation where the rows are sequences and the columns represent aligned symbols. Each column of an alignment must contain at least one symbol of Σ, in other words, a column with all gaps is not allowed. An example of multiple sequence alignment may be:

- Unaligned Sequences (input):

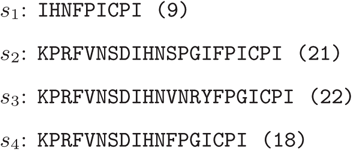
- Aligned Sequences (output):

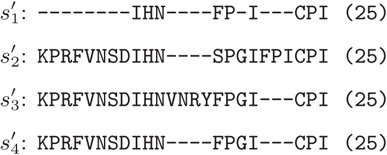

To find an accurate alignment, we propose the use of multiobjective optimization. Therefore, we search the best solution (alignment) that simultaneously maximizes the weighted sum-of-pairs function with affine gap penalties (WSP, *f*_1_) [9] and the number of Totally Conserved (*f*_2_) columns score [5], [21].

On the one hand, the weighted sum of pairs with affine gaps (WSP, *f*_1_) needs to maximize the following equation:

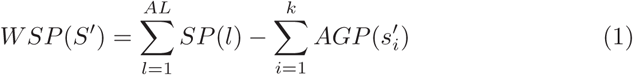

In equation 1, *AL* is the alignment length, *SP*(*l*) is the sum-of-pairs score of the *l^th^*column, which is defined as:

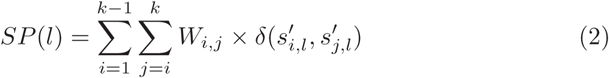

Note that in equation 2, *δ* is the substitution matrix used, either Pointed Accepted Mutation (PAM) or Block Substitution Matrix (BLOSUM); and *W_i,j_* refers to the sequence weight between sequence *s_i_* and *s_j_*. To compute the weight between two sequences we use the following equation:

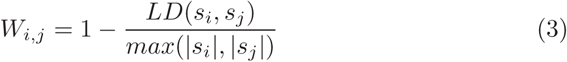

The Levenshtein Distance (*LD*) between two non-aligned sequences is the minimum number of *insertions, deletions* or *substitutions* required to change one sequence into the other.

In equation 1, *AGP* (*s*^′^_*i*_) is the a ne gap penalty score of sequence *s*^′^_*i*_:

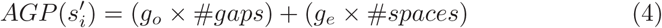

where *g_o_* is the weight to open the gap and *g_e_* is the weight to extend the gap with one more space. In this work, we have used the BLOSUM62 substitution matrix, *g_o_* = 6, and *g_e_* = 0.85.

On the other hand, the number of Totally Conserved (*f*_2_) columns score refers to the number of columns that are completely aligned with exactly the same compound. This objective function needs to be maximized to ensure more conserved regions within the alignment.

The maximum number of columns (alignment length) was limited to:

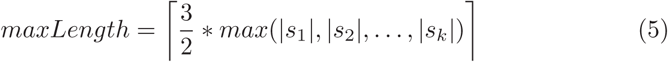

The choice of 1.5 as a scaling factor allowed the alignment to be 50% longer than the longest sequence in the set. This choice was based on the observation that solutions to common alignment problems rarely contained more than 50% gaps.

## 3 Parallel H4MSA

The Shuffled Frog-Leaping optimization Algorithm (SFLA) is a memetic meta-heuristic developed by Eusuff and Lansey [6]. The search procedure begins with a random population of frogs (solutions) in a swamp.

The population of frogs is divided into isolated communities (memeplexes) that will evolve independently, allowing different directions within the search space. At each community, the frogs evolve by sharing their ideas with their neighbour frogs. In this way, those frogs with better ideas will share more ideas than those frogs with poor ideas.

In addition, the best frogs of each community will share their ideas with other frogs in different communities. After a number of iterations, the communities of frogs are forced to mix and new communities are formed through a shu ing process in order to accelerate the convergence of the algorithm.

The chromosome representation of a solution in H4MSA is different from the traditional binary representation (’1’ indicates gap symbol and ‘0’ indicates a residue). For example, the binary representation of the alignment shown in Section 2 will be:

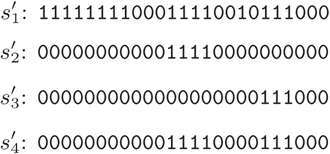

In H4MSA, a solution only stores the *number of groups of gaps* followed by the information of each group: *position of the first gap and number of successive gaps* (negative value). The chromosome representation of the example alignment will be:

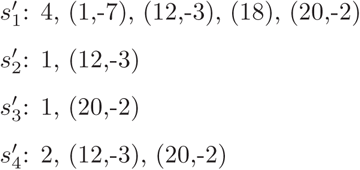

Given a set of *k* unaligned sequences (*S*), *m* memeplexes (number of communities) with *n* frogs per memeplex, a fixed number of evolutionary steps (*N*), and a stopping criterion, the procedure of H4MSA is:

1. Generate and evaluate *m* × *n* random alignments/frogs.
2. Sort the *m* × *n* frogs by alignment quality (*f*_1_) and conservation (*f*_2_). In this work, we have used the *Fast Non-Dominated Sorting* procedure [2].
3. Divide the *m* × *n* frogs into m memeplexes, such the first best frog goes to the first memeplex (*Y*_1_), the second best frog to the second memeplex (*Y*_2_), the *m^th^* best frog to the *m^th^* memeplex (*Y_m_*), the *m* + 1^*th*^ best frog to *Y*_1_, and so on.
4. For each memeplex (*Y_i_*)

a. Select the local worst (*X_lw_*) and local best frog (*X_lb_*) of the memeplex.
b. *X_lw_* learns from *X_lb_*, that is to say, *X_lw_* replaces a portion of its alignment with information obtained from *X_lb_*, generating a new frog (*X_new_*). For example, we select the following portion of the local best alignment:

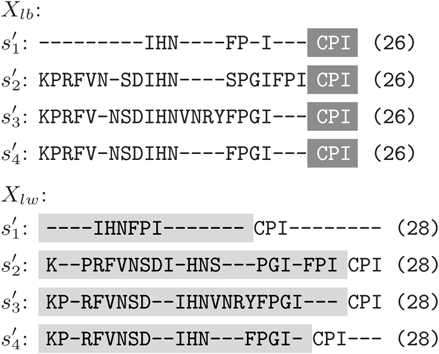

and the resultant new frog (*X_new_*) is constructed by taking into account both portions, filling with gaps until the length of all the sequences is exactly the same:

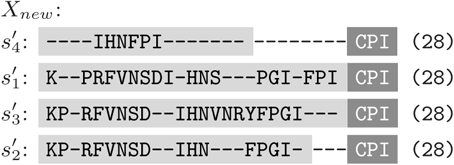
c. Apply the following mutation process to *X_new_*:

i. *Move a block*: randomly select a block of gaps/compounds and move it one position towards the left or right.

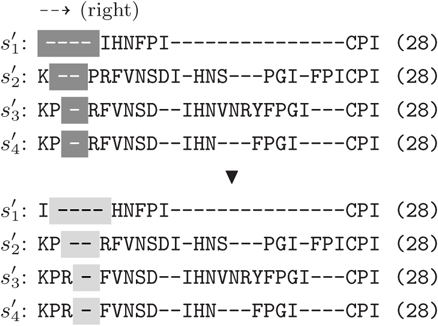
ii. *Merge two groups*: randomly select one of the sequences, choose a group of gaps/compounds, and merge it with the closest group.

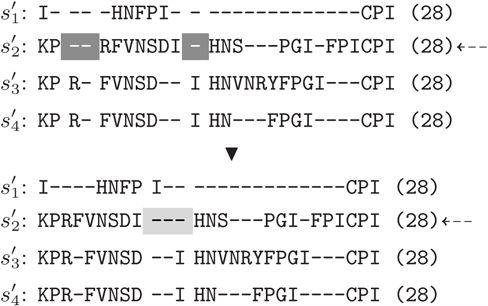
iii. *Divide a group*: randomly select one of the sequences, choose a group of gaps/compounds, and divided it into two new groups of approximately the same size.

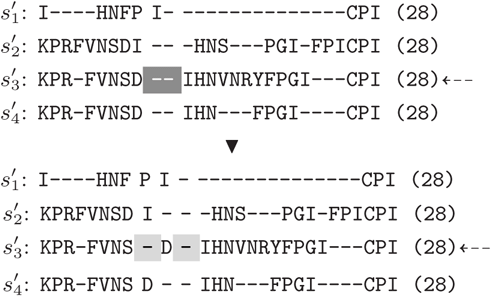
iv. *Compact Alignment*: delete those columns with all gaps.

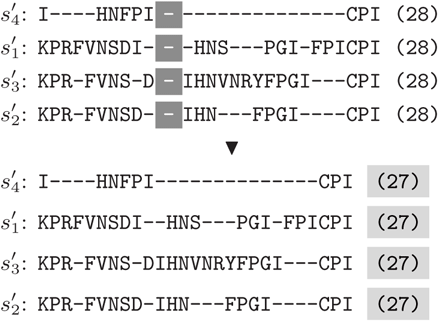
d. Evaluate *X_new_*, if *X_new_* is better than *X_lw_*, then go to Step 4(j).
e. Select the global best frog (*X_gb_*).
f. *X_lw_* learns from *X_gb_*, generating a new frog (*X_new_*).
g. Apply the mutation process to *X_new_*.
h. Evaluate *X_new_*, if *X_new_* is better than *X_lw_*, then go to Step 4(j).
i. Apply a *Local Search* to *X_lw_* and evaluate the new alignment produced. In our local search procedure, we use the fast and accurate Kalign2 [11] with the aim of re-aligning a portion of the input alignment. In the following, we present a step-by-step procedure of the local search:

i. Compute a random position of the alignment and a random size in the range [5-25%] of the alignment length:

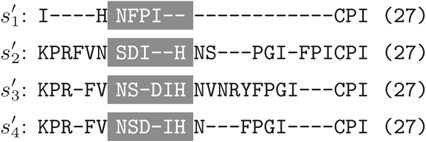
ii. Remove all gaps in the selected portion:

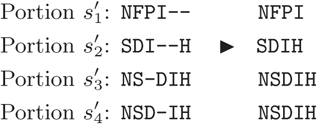
iii. Re-align the portion with Kalign2 [11] method. Input: Output:

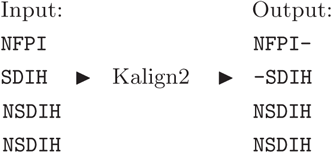
iv. The new portion re-aligned by Kalign2 replaces the old portion:

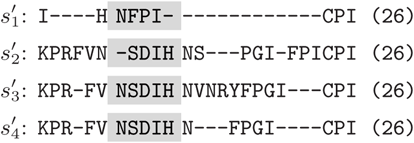
j. Replace *X_lw_* by *X_new_* and update the set of non-dominated solutions with *X_new_*.
k. If the maximum number of evolutionary steps (*N*) has not been reached, then go to Step 4(a). Otherwise, continue with the next memeplex.
5. Merge the frogs from the *m* memeplexes.
6. If the stopping criterion is satisfied, output the set of non-dominated solutions; otherwise, go to Step 2.

As we can see, the output of H4MSA is a set of non-dominated solutions, that is, a set of solutions that represents a trade-off between alignment quality (*f*_1_) and consistency (*f*_2_). A detailed description of the H4MSA algorithms appears in [18].

In this work, we propose a parallel scheme of H4MSA for a shared-memory architecture. After studying the computational requirements of H4MSA, we can see that a large amount of time is spent in the initial generation of the population and in the evolving process of each memeplex; therefore, we focus on parallelizing these tasks.

On the one hand, in the generation of the initial population (Step 1), the *m* × *n* random alignments are divided among the available threads. So, if the number of threads is equal to *M*, each thread will be in charge of generating and evaluating 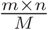 random alignments (frogs). In Figure 1, we can see an illustrative comparison between the sequential and parallel procedure.

**Figure 1:**
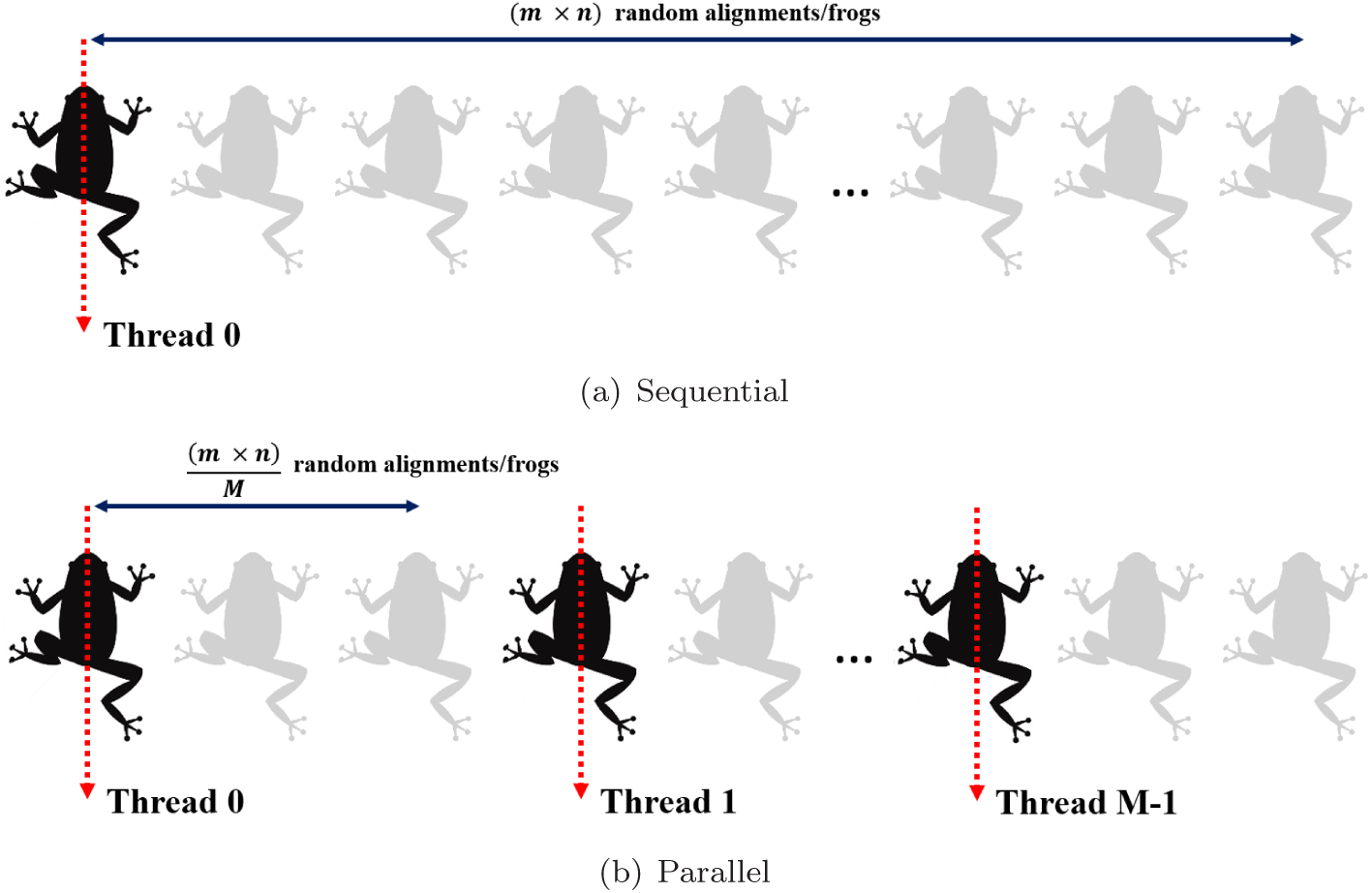
Sequential vs. Parallel generation of the initial population in H4MSA.

The second and third steps of H4MSA are carried out only by one thread, because the sorting and division process consumes insignificant runtime in comparison with other tasks.

In the fourth step, H4MSA performs m independent evolutionary processes of N iterations. As we can see, the tasks carried out within each iteration of the evolutionary process are computationally expensive: learning process, mutation process, and local search procedure. This loop has been parallelized, but we have defined a critical section when the *X_lw_* is replaced by the *X_new_* and the set of non-dominated solutions is updated (Step 4(j)).

In the evolving process loop, the workload of each iteration may be di erent for each thread; therefore, the distribution of iterations is not static, that is, each thread is in charge of performing the consecutive 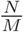 iterations of the evolutionary process, assuming *M* available threads. On the contrary, in this approach, we use a dynamic distribution scheduling of the *N* iterations among the threads during runtime.

Since H4MSA has been implemented in OpenMP, we have parallelized the evolving of each memeplex by using the `dynamic schedule`. This schedule uses an internal work queue with the loop iterations to each thread. When a thread is finished, it retrieves the next loop iteration from the top of the work queue. The number of iterations performed by each thread is decided during the execution of the algorithm; so, threads may do different number of iterations. In Figure 2, we show the advantages of using a dynamic distribution of the iterations among threads when the workload of iterations is different. As we can see, 51.51% of efficiency is achieved with an static schedule; however, the efficiency increases up to 94.45% when the distribution of the iterations is dynamic.

**Figure 2:**
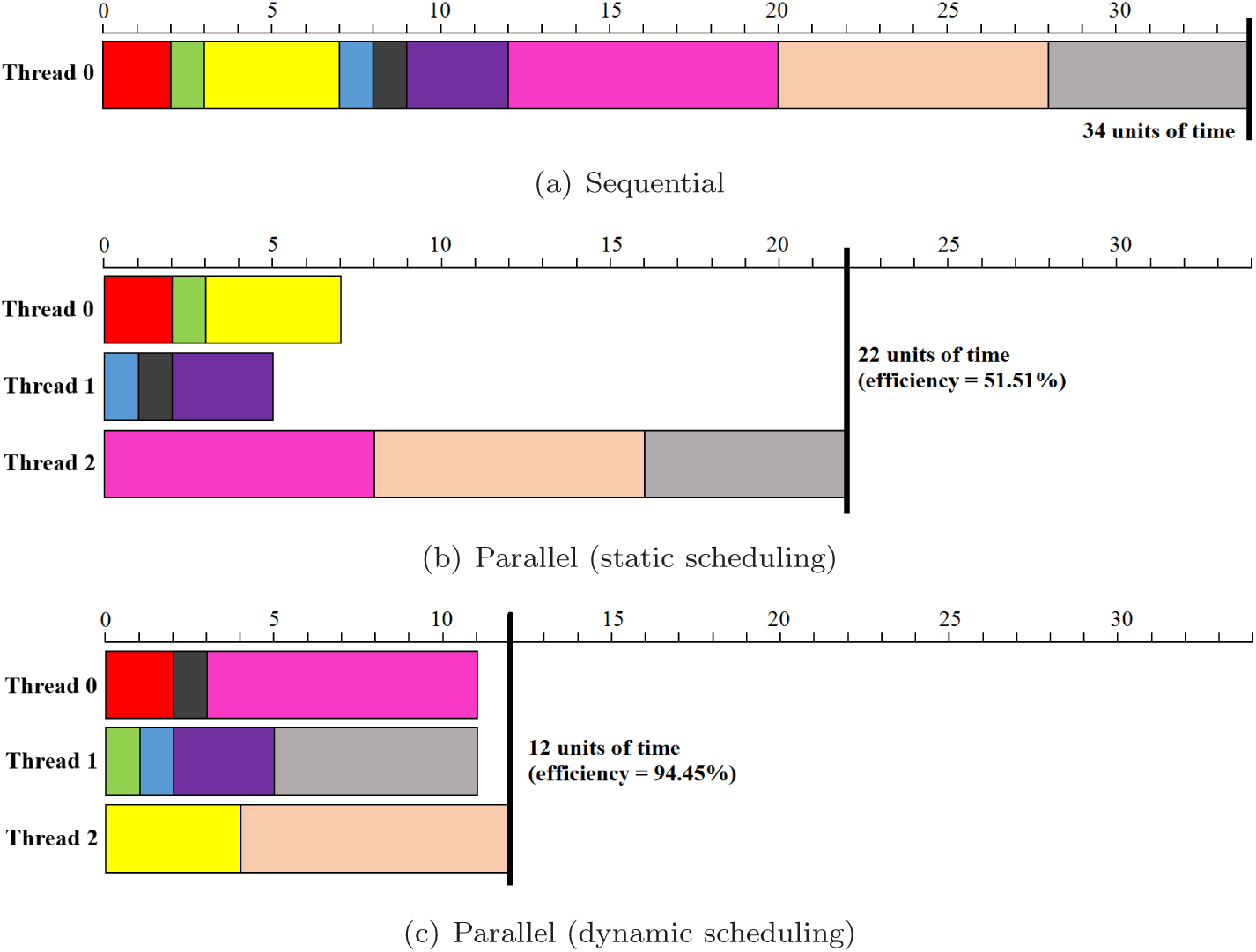
Static vs. dynamic distribution of iterations.

## 4 Experimental Results

In this section we present a comparative study between our parallel approach (H4MSA) and other multi-threaded MSA approaches published in the literature.

We have compared the multithreaded version of H4MSA with those multiple sequence alignment approaches that allow being run in multi-core environments:

- **MSAProbs** (version 0.9.7) [12]. It is a parallel and accurate approach for MSA. It allows the use of the -`num_threads` flag to specify the number of threads.
- **T-Coffee** (version 11.00.8) [15]. It is a widely used MSA approach in the field. The steps of T-Coffee are multi-threaded by using the `-multi_core` flag, specifying the number of cores to use by the `-n_core` flag.
- **Clustal** Ω (version 1.2.1) [19]. It is the latest addition to the Clustal family. It offers a significant increase in scalability over previous versions, allowing a large number of sequences to be aligned. It also make use of multiple processors by using the `--threads` flag.
- **MAFFT** (version 7.215) [10]. It is a method for rapid multiple sequence alignment based on fast Fourier transform. From version 6.8, MAFFT switches to the multi-core version by simply specifying the number of threads with the `--thread` flag.

The aforementioned approaches were run by using the default parameter configuration.

In H4MSA we found three main parameters: number of memeplexes (*m*), number of frogs at each memeplex (*n*), and the number of evolutionary steps (*N*). In this comparative study, we have used: *m* = 4 (4 memeplexes), *n* = 32 (32 frogs per memeplex), and *N* = 10 (10 evolutionary steps). The stopping criterion is based on the number of fitness evaluations: 50000 evaluations.

The datasets used in this experiments were taken from the HOMFAM benchmark suite [19]. As we can see in Table 1, we have chosen datasets with different number of sequences, from 88 sequences to 1056 sequences, in order to evaluate the performance of the approaches when the number of sequences increases.

**Table 1:**
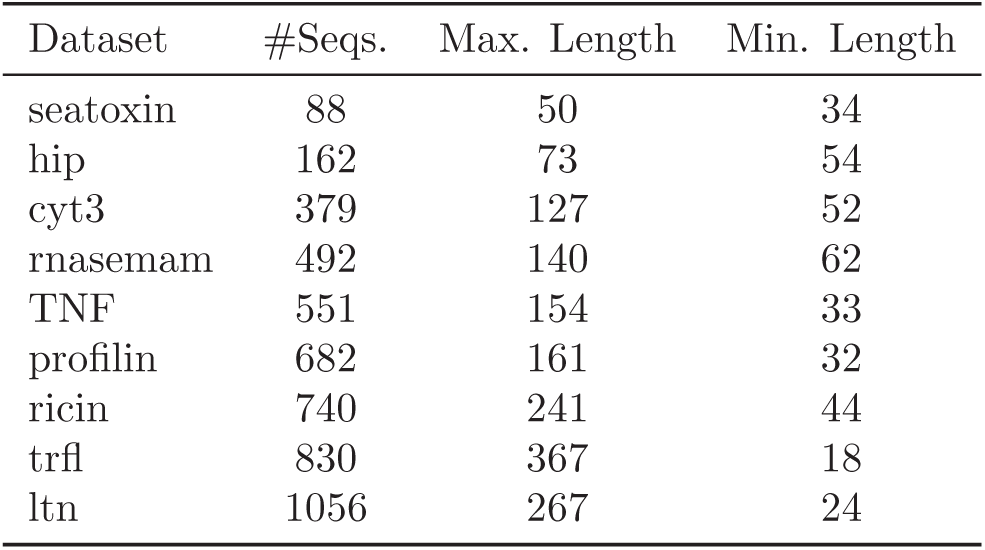
Selected HOMFAM datasets.

In order to extract useful conclusions with a certain level of statistical confidence, 30 independent runs were performed for each approach involved, and the average runtime was used. The architecture selected for conducting these experiments was a 2-processor AMD Opteron^TM^ Processor 6376 of 16 cores at 2.3GHz and 12MB Cache running Scientific Linux 6.1.

For measuring the performance of the parallel approaches, we have used two well-known metrics: *speedup*s and *eFFciency*. Amdahl’s Law states that the performance improvement to be gained from using some faster mode of execution is limited by the fraction of the time the faster mode can be used. Therefore, Amdahl’s Law defines the *speedup* that can be gained by using a particular feature. In a more formal way, let *T_M_* be the runtime for an algorithm using *M* threads and *T*_1_ the runtime of the sequential version, the speedup reports us how much faster an algorithm will run as opposed to the sequential version. The efficiency is computed dividing the obtained speedup with *M* threads by the number of threads used (*M*).

In order to determine the accuracy of each MSA method, we have evaluated the level of conservation with the BLOSUM62 substitution matrix. Therefore, the alignment obtained by each approach for each data set was scored by using the `+evaluate blosum62mt` action provided by T-Coffee. Note that, a higher score implies better alignment accuracy.

On the one hand, in Table 2, we present the conservation score obtained by H4MSA, MSAProbs, T-Coffee, Clustal Ω, and MAFFT. In Figure 3, we present a visual comparison among the five approaches in terms of conservation score. As we can see in Figure 3, the alignment accuracies obtained by H4MSA in all the datasets tested are better than the well-known approaches. In addition, if we focus on the largest dataset (*ltn*, 1056 sequences), we observe an average conservation improvement around 14.98%. Therefore, we can conclude that H4MSA is able to obtain accurate alignments.

**Table 2:** Average conservation score in 30 independent runs.

| Dataset | H4MSA | MSAProbs | T-Coffee | Clustal $\Omega$ | MAFFT |
| --- | --- | --- | --- | --- | --- |
| seatoxin | **704** | 688 | 686 | 681 | 689 |
| hip | **605** | 590 | 595 | 592 | 594 |
| cyt3 | **485** | 347 | 357 | 367 | 389 |
| rnasemam | **572** | 547 | 550 | 546 | 550 |
| TNF | **527** | 462 | 466 | 463 | 467 |
| profilin | **604** | 543 | 554 | 541 | 551 |
| ricin | **576** | 472 | 493 | 477 | 485 |
| trfl | **588** | 537 | 542 | 544 | 546 |
| ltn | **584** | 485 | 501 | 499 | 501 |

**Figure 3:**
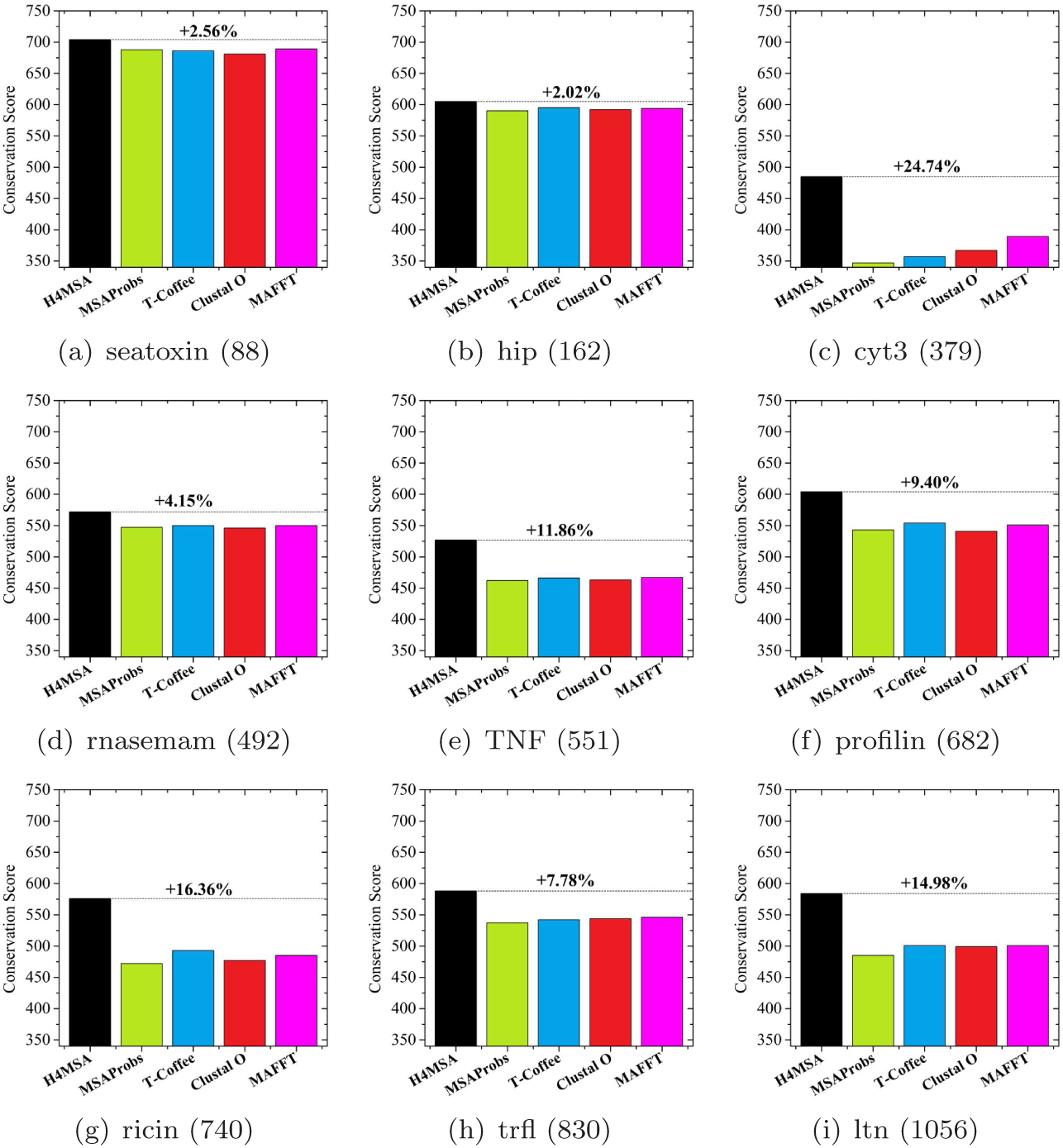
Comparison among H4MSA, MSAProbs, T-Coffee, Clustal Ω, and MAFFT in terms of average Conservation Score. Note that, the number of sequences appears between brackets for each dataset.

On the other hand, we compare the parallel performance of the approaches under study. In Table 3 and Table 4, we present the sequential runtime of each method and the parallel speedup and efficiency obtained with different number of cores (2, 4, 8, 16, and 32 cores); respectively. As we can see, the parallel performance of Clustal Ω, T-Coffee, and MAFFT is very poor when the number of cores increases. We can also observe that MSAProbs presents a nice parallel performance with 2, 4, and 8 cores, but its efficiency decreases with 16 and 32 cores. The parallel efficiency of H4MSA remains over 75% in all cases. In Figure 4 and 5, we present a comparison among the five parallel approaches in terms of runtime, speedup, and efficiency.

**Figure 4:**
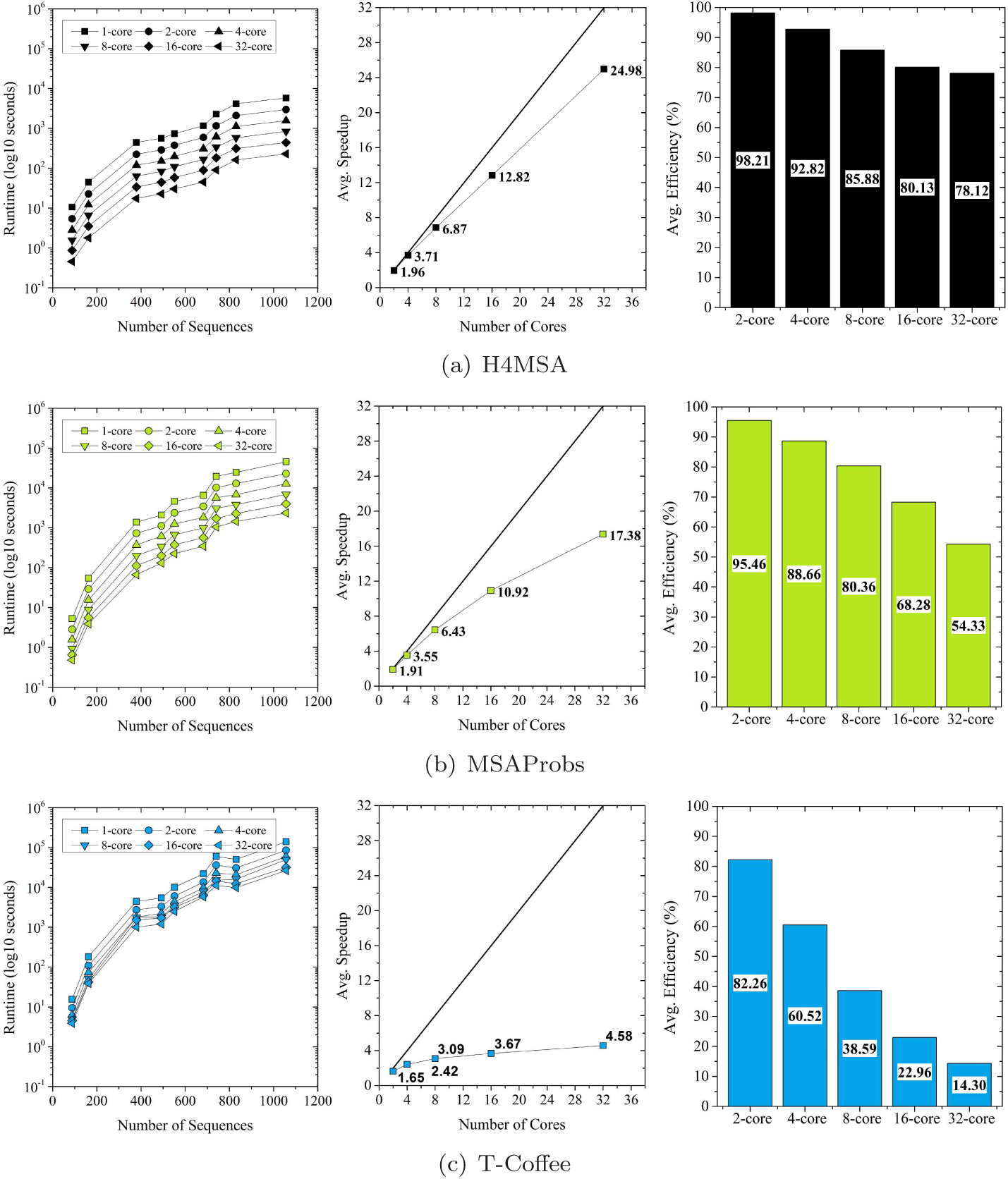
Runtime, average speedup, and average efficiency (%) obtained by each MSA tool.

**Figure 5:**
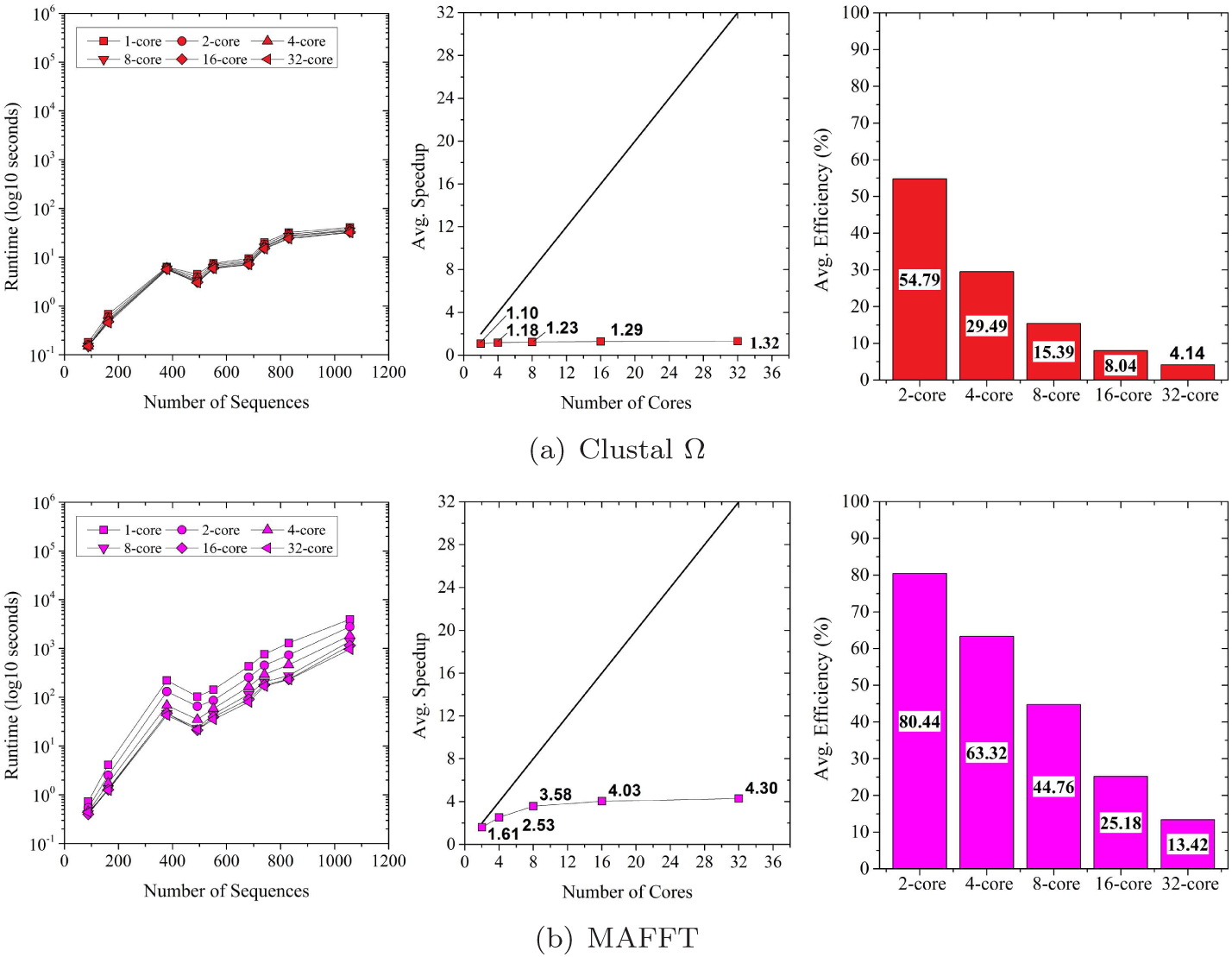
Runtime, average speedup, and average efficiency (%) obtained by each MSA tool.

**Table 3:**
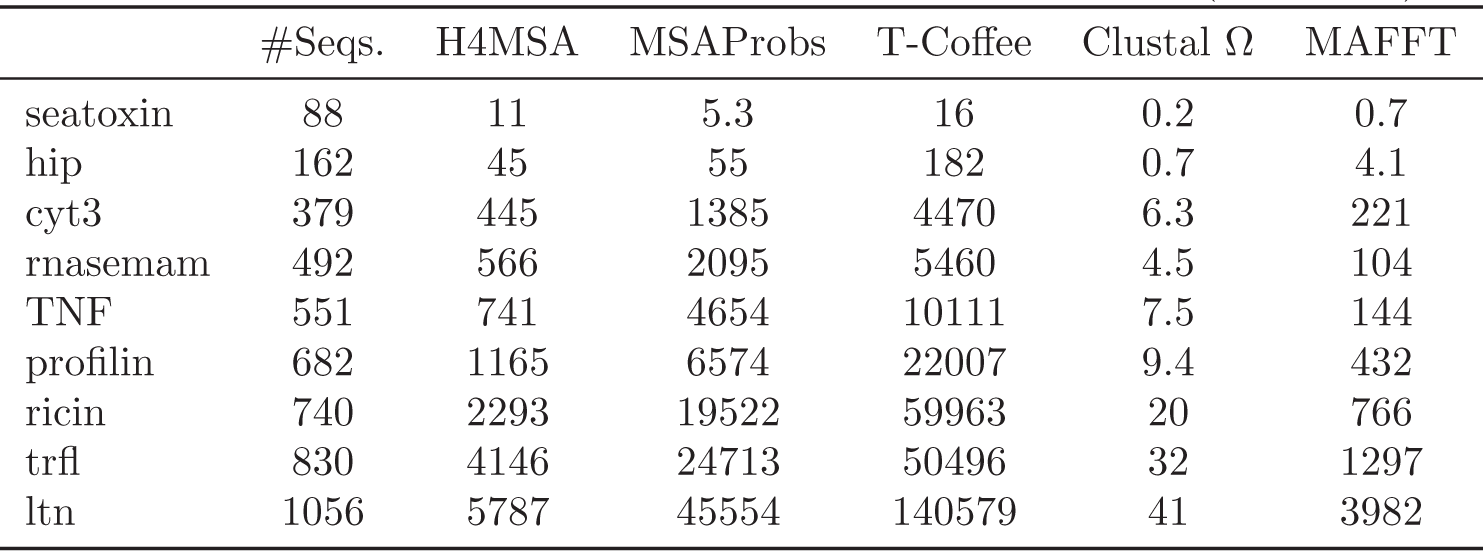
Runtime spent by the sequential version of each aligner (in seconds)

**Table 4:** Speedup and Effciency (%) obtained by the multi-threaded MSA approaches.

| | H4MSA | | MSAProbs | | T-Coffee | | Clustal $\Omega$ | | MAFFT | |
| --- | --- | --- | --- | --- | --- | --- | --- | --- | --- | --- |
| #Cores | *S* | *E* | *S* | *E* | *S* | *E* | *S* | *E* | *S* | *E* |
| 2 | 1.96 | 98.21 | 1.91 | 95.46 | 1.65 | 82.26 | 1.10 | 54.79 | 1.61 | 80.44 |
| 4 | 3.71 | 92.82 | 3.55 | 88.66 | 2.42 | 60.52 | 1.18 | 29.49 | 2.53 | 63.32 |
| 8 | 6.87 | 85.88 | 6.43 | 80.36 | 3.09 | 38.59 | 1.23 | 15.39 | 3.58 | 44.76 |
| 16 | 12.82 | 80.13 | 10.92 | 68.28 | 3.67 | 22.96 | 1.29 | 8.04 | 4.03 | 25.18 |
| 32 | 24.98 | 78.12 | 17.38 | 54.33 | 4.58 | 14.30 | 1.32 | 4.14 | 4.30 | 13.42 |

*S*: Average speedup in the nine datasets of HOMFAM.
*E*: Average efficiency (%) in the nine datasets of HOMFAM.

In Figure 4 and 5, if we compare the sequential runtime of each approach, we can see that the fastest algorithms are (in order): Clustal Ω, MAFFT, H4MSA, MSAProbs, and T-Coffee. However, thanks to the parallel efficiency of H4MSA, it is able to be the second fastest approach when the number of cores is set to 32.

All in all, we can conclude that H4MSA is not only an accurate alignment method but also its parallel performance allows it to handle datasets with hundreds of sequences in a reasonable amount of time.

## 5 Conclusions and Future Works

A parallelization of the Hybrid Multiobjective Memetic Metaheuristic for Multiple Sequence Alignment (H4MSA) is presented in this work. H4MSA is based on the Shuffled Frog-Leaping Algorithm, which provides the benefits of information mixture of the ‘shuffled complex evolution’ technique. The parallel version of H4MSA has been compared with the parallel approaches of MSAProbs, T-Coffee, Clustal Ω, and MAFFT when solving datasets with different number of sequences in the range [88–1056]. We can conclude that the alignment accuracy and parallel performance of H4MSA is significantly better than other approaches published in the literature.

As future work, we intend to develop a parallel version of H4MSA for shared-and distributed-memory architectures, in order to solve larger datasets.

## Acknowledgements

Álvaro Rubio-Largo is supported by the post-doctoral fellowship SFRH/BPD/100872/2014 granted by Fundação para a Ciência e a Tecnologia (FCT), Portugal.

